# Reconstruction of the cell entry pathway of an extinct virus

**DOI:** 10.1101/331363

**Authors:** Lindsey R. Robinson-McCarthy, Kevin R. McCarthy, Matthijs Raaben, Silvia Piccinotti, Joppe Nieuwenhuis, Sarah H. Stubbs, Mark J.G. Bakkers, Sean P. J. Whelan

## Abstract

Endogenous retroviruses (ERVs), remnants of ancient germline infections, comprise 8% of the human genome. The most recently integrated includes human ERV-K (HERV-K) where several envelope (*env*) sequences remain intact. Viral pseudotypes decorated with one of those Envs are infectious. Using a recombinant vesicular stomatitis virus encoding HERV-K Env as its sole attachment and fusion protein (VSV-HERVK) we conducted a genome-wide haploid genetic screen to interrogate the host requirements for infection. This screen identified 11 genes involved in heparan sulfate biosynthesis. Genetic inhibition or chemical removal of heparan sulfate and addition of excess soluble heparan sulfate inhibit infection. Direct binding of heparin to soluble HERVK Env and purified VSV-HERVK defines it as critical for viral attachment. Cell surface bound VSV-HERVK particles are triggered to infect on exposure to acidic-pH, whereas acid pH pretreatment of virions blocks infection. Testing of additional endogenous HERV-K *env* sequences reveals they bind heparin and mediate acid pH triggered fusion. This work reconstructs and defines key steps in the infectious entry pathway of an extinct virus.

**Author Summary:** The genomes of all vertebrates are littered with the remains of once exogenous retroviruses. The properties of these ancient viruses that fostered germline colonization and their subsequent inheritance as genetic elements are largely unknown. The viral envelope protein (Env) dictates the cell entry pathway. Here we define host factors involved in the cell-entry of the youngest human ERV, HERV-K. Using a forward genetic screen, we identified heparan sulfate as a critical mediator of productive cell-entry. The abundance of this carbohydrate on almost all cells in the body suggests that HERV-K endogenization was a consequence of a broad tropism and not a specific targeting of germ cells. We demonstrate that multiple HERV-K Env protein encoded in the genome bind heparin. As HERV-K Envs are expressed in some transformed and virus-infected cells as well as during inflammation, it is tempting to speculate that this heparan sulfate binding property could be physiologically relevant during disease.

## Introduction

Endogenous retroviruses (ERVs) are remnants of ancient germline infections and comprise approximately 8% of the human genome [1]. The degraded nature of ERV sequences impedes investigation of the properties of the infectious progenitor viruses and the events that led to their endogenization. During evolution, ERV sequences accumulate mutations, consequently the most recently endogenized sequences are the most likely to reflect the properties of the progenitor virus from which they were derived. The most recent human endogenous retroviruses (HERVs) belong to the HERV-K (HML-2) group. Multiple endogenization events resulted in approximately 90 proviral copies and 1,000 solo long terminal repeats (LTRs) in the reference human genome [2]. The HERV-K (HML-2) group is approximately 30-35 million years old [3], with evidence of endogenization as recently as 100,000-600,000 years ago [4, 5].

Many HERV-K sequences exist as largely intact proviral copies, some of which still encode single functional proteins [6]. While no single locus has been demonstrated to produce an infectious virus, many loci have retained the capacity to produce individual functional proteins. For example, at least one copy, termed HERV-K 108, has retained the capacity to produce an envelope (Env) that can mediate cellular attachment and entry [7]. Two replication-competent infectious clones, Phoenix [8] and HERV-K_CON_ [9] have been reconstructed from consensus sequences comprising the most recently endogenized loci. The reconstructed viruses grow poorly which has hampered efforts to study the biology of their envelope proteins.

The processes that govern endogenization are poorly defined. The first virus-cell contacts are mediated through viral glycoproteins, which can dictate species, tissue and cellular tropism. We have previously overcome some of the challenges imposed by viral titer by generating an infectious vesicular stomatitis virus (VSV) in which the glycoprotein was replaced by Phoenix Env (VSV-HERVK). Using this virus we determined that HERV-K Env imparts a broad species and tissue tropism [10] and demonstrate that productive infection of mammalian cells requires access to an acidified compartment that is accessed via a dynamin-dependent but clathrin-independent pathway [10]. We also found that proteolytic processing and acid pH are required for HERV-K Env to mediate membrane fusion. A broad species and cell-type tropism was also described for a modified variant of a different ancestral sequence [11]. The broad host range reported in those studies implies that host factors required for HERV-K entry are evolutionarily conserved and ubiquitously expressed.

To identify such host factors we performed a genome-wide haploid genetic screen by selecting cells resistant to VSV-HERVK infection. This approach has identified critical host factors required for the entry of several extant viruses, including Ebola, Lassa, Lujo, Andes virus, and Rift Valley fever virus [12-17]. We identify genes involved in heparan sulfate biosynthesis and demonstrate a specific interaction between this glycosaminoglycan and multiple HERV-K envelope proteins. We further show that acid pH is required to trigger membrane fusion by these Envs and is sufficient to mediate infection of cell surface virus and to inactivate unbound virions. Based on our findings we posit a model for the entry pathway of this extinct virus where heparan sulfate binding followed by subsequent endosomal uptake and acidification result in productive infection.

## Results

To identify host factors required for HERV-K Env mediated entry, we performed a haploid genetic screen [12] (Fig 1A-B). Briefly, HAP1 cells were mutagenized using a retroviral gene-trap vector to generate a population with inactivating mutations across the genome and infected with VSV-HERVK. Deep sequencing genomic DNA from cells that survived VSV-HERVK infection identified sites of integration of the gene-trap retrovirus (Fig 1C). Among the genes identified were 11 involved in the biosynthesis of heparan sulfate - a glycosaminoglycan (GAG) ubiquitously expressed on the cell surface. Six of those genes (*GPC3, EXT1, EXT2, EXTL3, HS2ST1,* and *NDST1*) are specific to heparan sulfate and heparin and not other GAGs (S1 Fig). For follow up, we selected *EXT1* which encodes an enzyme that catalyzes the addition of a glucaronic acid – N-aceytlglucosamine (GlcA-sGlcNAc) disaccharide onto the growing heparan sulfate chain and *SLC35B2* which encodes the Golgi-resident transporter of the universal sulfate donor 3’-phosphoadenosine-5’-phosphosulfate (PAPS) [18]. Three additional genes, myosin X (*MYO10)*, sortilin (*SORT1*), and CREB binding protein (*CREBBP*) scored as significant and were also selected for further follow up.

**Fig 1.**
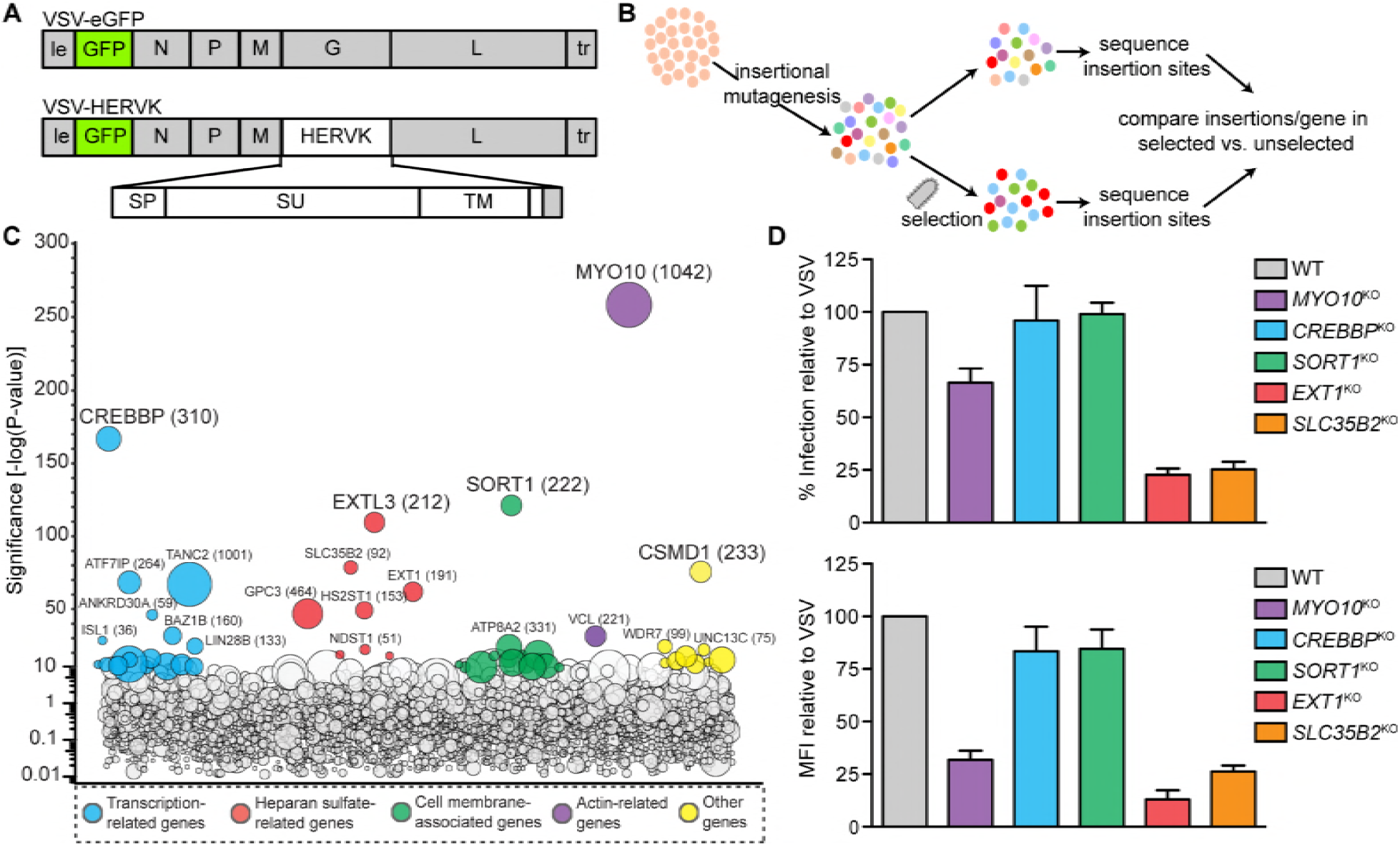
Haploid genetic screen identifies host factors required for infection by VSV-HERVK. (**A)** Viral genome structures. Sequences from VSV are shown in grey: N, nucleocapsid; P, phosphoprotein; M, matrix; G, glycoprotein; L, large polymerase. le: leader. tr: trailer. Viruses also encode an eGFP reporter gene. VSV-HERVK encodes the HERVK glycoprotein, which contains the signal peptide (SP), surface (SU) subunit, transmembrane subunit (TM), and membrane-spanning domain of HERV-K env, and cytoplasmic tail of VSV G. **(B)** Schematic of haploid genetic screen. HAP1 cells were subjected to insertional mutagenesis, followed by selection with VSV-HERVK. Surviving cells were deep sequenced to identify the position of insertion sites. The number of insertions per gene in the selected set was compared to that of an unselected set to identify genes that were associated with survival of infection. **(C)** Screen results. The y-axis indicates the significance of enrichment of gene-trap insertions compared with unselected control cells. Circles represent individual genes and their size corresponds to the number of unique insertion sites in the selected population. Genes with significance scores above 10 are colored according to function and grouped horizontally. Genes with significance score above 25 are labeled. **(D)** HAP1 cells were gene edited to lack the indicated genes and infected with VSV or VSV-HERVK. The fold difference in percent infected cells (top) and mean fluorescence intensity (MFI, bottom) of VSV-HERVK infected cells normalized to that of VSV is shown. Error bars represent standard error of the mean (SEM) for at least three independent experiments.

We independently generated single-cell clones of HAP1 cells lacking each of those 5 genes by gene editing and infected them with VSV or VSV-HERVK expressing eGFP as a marker of infection (Fig 1D). This eliminated *MYO10*, *SORT1* and *CREBBP* from further analysis in viral entry because the fraction of cells infected was only modestly changed (Fig 1D and S2 Fig). In *MYO10*^KO^ cells we note however that the intensity of eGFP expression increased following infection with VSV but slightly decreased following infection with VSV-HERVK (S2 Fig). This result indicates that elimination of myosin X differentially impacts the kinetics of productive infection perhaps reflective of the distinct uptake mechanisms of VSV compared to VSV-HERVK (S2 Fig). By contrast, VSV-HERVK infection of *EXT1*^*KO*^ and *SLC35B2*^*KO*^ cells was reduced 4-fold compared to VSV. VSV infection was unaffected in *EXT1*^*KO*^ cells, but was diminished 3-fold in *SLC35B2*^*KO*^ cells. Those results demonstrate that elimination of cell surface heparan sulfate reduces VSV-HERVK infection specifically and that suppression of sulfation also reduces VSV infection in a manner that appears independent of heparan sulfate (S2 Fig). Flow cytometry verified that cell surface expression of heparan sulfate was lost in both *EXT1*^*KO*^ and *SLC35B2*^*KO*^ cells and was restored following transduction with retroviruses expressing the corresponding gene (S3 Fig). Restoration of cell surface heparan sulfate corresponded with an increase in VSV-HERVK infection (Fig 2A). Loss of heparan sulfate did not completely block VSV-HERVK infection as evident from the small fraction of infected cells. That small fraction, however, exhibits a 2-3 fold reduction in the intensity of eGFP, presumably reflecting a less efficient heparan sulfate independent mechanism of viral entry (Fig 2B and S4 Fig).

**Fig 2.**
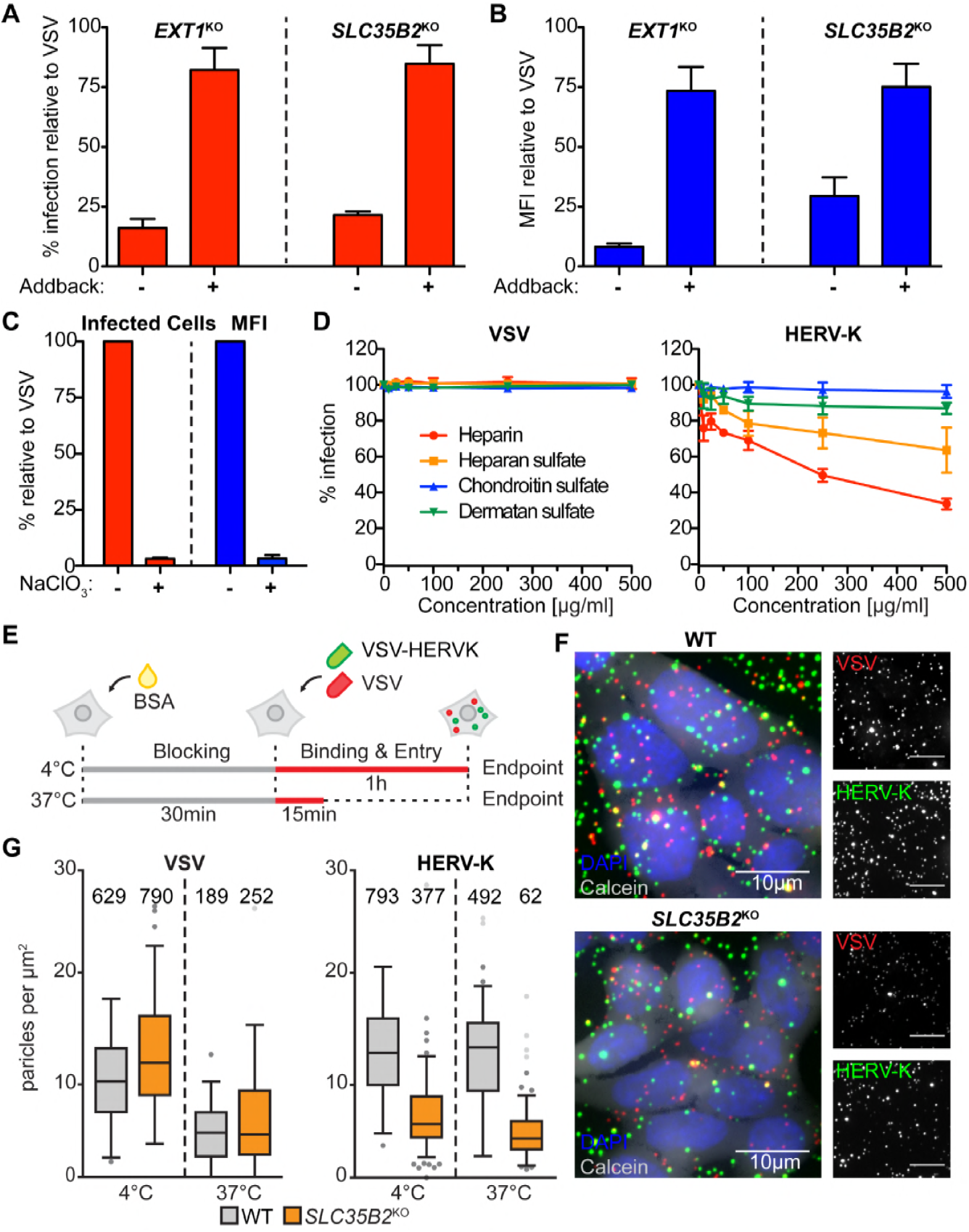
Heparan sulfate facilitates HERV-K Env-mediated entry and attachment. (A) WT HAP1, *EXT1*^*KO*^ Neo^r^ Ctrl (transduced with a control retrovirus), *EXT1*^*KO*^ EXT1-HA (+ addback), *SLC35B2*^*KO*^ Neo^r^ Ctrl and *SLC35B2*^*KO*^ SLC25B2-HA (+ addback) cells were infected with VSV-HERVK or VSV and infectivity analyzed by flow cytometry. Fold difference in percent infected cells compared to WT for VSV-HERVK was normalized to that of VSV for each condition. Error bars represent SEM for four independent experiments. **(B)** Relative MFI of cells from (A). Data were normalized as in (A). **(C)** BSRT7 cells were treated with 50mM sodium chlorate and infected with either VSV or VSV-HERVK. Fold difference in both percent infected cells (left) and MFI (right) compared to untreated cells for VSV-HERVK was normalized to that of VSV. Error bars represent SEM for three independent experiments. **(D)** VSV or VSV-HERVK was incubated with the indicated soluble glycosaminoglycans prior to infecting BSRT7 cells. Percent infected cells was normalized to untreated virus controls. Error bars represent SEM for three independent experiments. **(E)** Schematic of virus attachment experiment. Cells were blocked with BSA then incubated with both fluorescently labeled VSV-HERVK and VSV at either 37°C or 4°C. **(F)** Representative images from 4°C attachment experiment. Red: VSV. Green: VSV-HERVK. Blue: DAPI. Grey: calcein. **(G)** Results of attachment experiment. Numbers of particles/μm^2^ are plotted. Grey circles indicate outliers. Total number of particles counted per condition is indicated above each box.

As a complementary approach to genetic inactivation of heparan sulfate biosynthesis, we employed a chemical approach. Sodium chlorate treatment of cells inhibits the synthesis of PAPS and correspondingly reduces cell surface sulfation. Cells cultured in the presence of 50 mM sodium chlorate showed a 30-fold reduction in infectivity of VSV-HERVK compared to VSV (Fig 2C). The fraction of cells that were infected by VSV-HERVK again showed a reduction in the levels of eGFP expressed, following entry independent of heparan sulfate (Fig 2C and S4 Fig). These results confirm the findings obtained following genetic inactivation of heparan sulfate biosynthesis and support a role for heparan sulfate in HERV-K entry.

Heparan sulfate has been identified as a receptor for herpes simplex virus 1 (HSV1) [19] and eastern equine encephalitis virus (EEEV) [20]. If heparan sulfate serves as a key entry factor for HERV-K, VSV-HERVK infection should be sensitive to competition by excess soluble GAGs. Incubation of purified virus with soluble heparin - a highly sulfated analog of heparan sulfate - or with heparan sulfate, inhibits infection in a concentration dependent manner. The sulfated GAGs chondroitin or dermatan sulfate had no effect on VSV or VSV-HERVK infection further supporting a specific requirement for heparan sulfate in HERVK infection at the level of viral attachment (Fig 2D). Consistent with this interpretation, attachment of single VSV-HERVK particles to *SLC35B2*^*KO*^ cells was reduced at least 2-fold compared to WT cells (Fig 2E-G) at both 4°C and 37°C. By contrast VSV particle binding was similar between both cell types at both temperatures (Fig 2G).

Further evidence that heparan sulfate serves an entry factor was provided by the demonstration that VSV-HERVK particles specifically associate with heparin but not protein A beads (Fig 3A). This heparin bead binding was sensitive to inhibition by pre-incubation of virus with soluble heparin. VSV did not bind either heparin or protein A beads, underscoring that binding is dictated by the HERV-K glycoprotein. To test whether HERV-K Env directly interacts with heparin, we generated a soluble, monomeric HERV-K SU subunit, which by extrapolation from extant retroviruses would harbor the receptor-binding domain (Fig 3B, S5 Fig, and S6 Fig). Soluble HERV-K SU specifically bound to heparin but not protein A beads, and this binding was sensitive to pre-incubation of the protein with soluble heparin (Fig 3C). As expected a soluble receptor-binding domain from Influenza A hemagglutinin (HA), which binds a different carbohydrate receptor, sialic acid, failed to bind either the heparin or protein A beads, further supporting the specificity of the HERVK-heparan sulfate interaction. Pre-incubation of HERV-K SU with soluble GAGs prior to mixing with heparin beads demonstrates that binding is inhibited by soluble heparin and heparan sulfate, but not chondroitin or dermatan sulfate (Fig 3D). These data correlate with the suppression of infectivity, and provide further evidence that heparan sulfate binding leads to productive infection by VSV-HERVK. We further found that HERV-K SU binding to heparin beads was unaffected by pre-incubation with 2-O-desulfated heparin, whereas 6-O-desulfated heparin showed partial inhibition of binding (Fig 3D). This result implies that 2-O sulfation and not 6-O sulfation is important for HERV-K binding. Consistent with this, our genetic screen identified the enzyme that catalyzes 2-O-sulfation (heparan sulfate 2-O-sulfotransferase 1 (HS2ST1)) but not the enzymes that catalyze 6-O-sulfation (HS6ST1,2,3).

**Fig 3.**
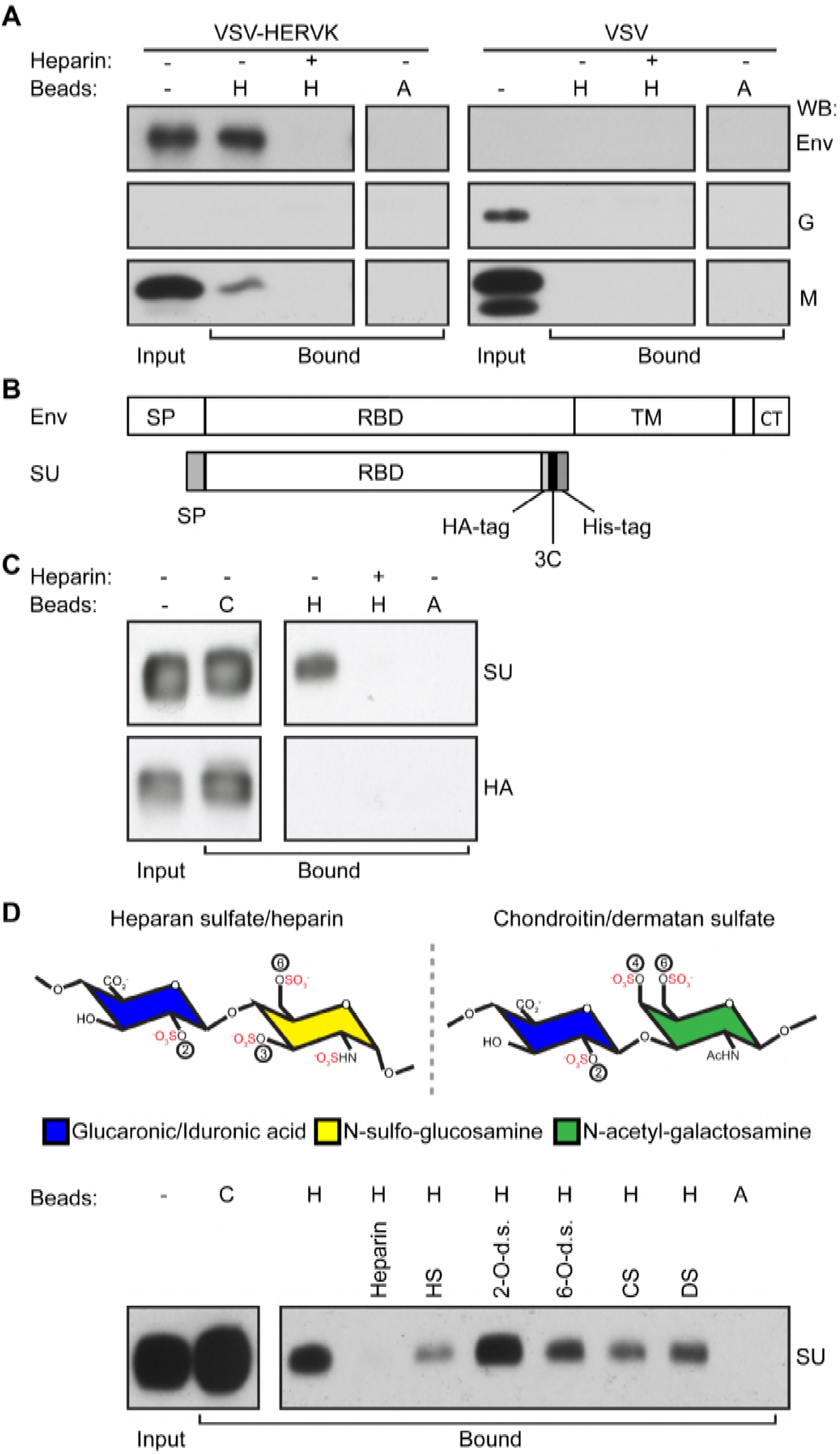
HERV-K Env binds heparin. (A) Purified VSV-HERVK or VSV were incubated with heparin (H) or protein A (A) beads, with (+) or without (-) soluble heparin added as a competitor. Bound virions were analyzed by Western blot against HERV-K Env, VSV G, and VSV M. Input: total input virus. **(B)** Schematic of HERV-K Env and HERV-K SU used in this study. SP: signal peptide. RBD: receptor binding domain. TM: transmembrane subunit. CT: cytoplasmic tail. 3C: 3C protease cleavage site. **(C)** HERV-K SU or Influenza A HA receptor binding domain (HA) were pre-incubated with or without soluble heparin prior to incubation with either cobalt (C, maximum pull-down control), heparin (H), or protein A (A) agarose beads. Bound protein was eluted from the beads and subjected to SDS-PAGE followed by Western blot against the HA tag. Input: 10% of total input protein. **(D)** Top: Structure of glycosaminoglycans. The repeating disaccharides of heparan sulfate/heparin (left) and chondroitin/dermatan sulfate (right) are shown. Sulfates are highlighted in red. Positions of O-sulfations are indicated with circled numbers. Disaccharides are shown as fully sulfated, however individual sugars will not always be sulfated at each position. Bottom: HERV-K SU was pre-incubated with soluble competitor compounds (heparin, heparan sulfate, 2-O-desulfated heparin, 6-O-desulfated heparin, chondroitin sulfate A, and dermatan sulfate) prior to incubation with either cobalt, heparin, or protein A agarose beads. Bound protein was eluted from the beads and subjected to SDS-PAGE followed by Western blot against the HA tag. Input: 10% of total input.

Acidic pH – such as that encountered in endocytic compartments – serves as the trigger for conformational rearrangements in several viral envelope proteins necessary for membrane fusion. In class I fusion proteins, such as influenza HA, those rearrangements are irreversible, such that premature exposure to acidic pH inactivates the fusion machinery. Envelope proteins from every extant betaretrovirus that has been tested, including Jaagsiekte sheep retrovirus (JSRV), enzootic nasal tumor virus (ENTV), and mouse mammary tumor virus (MMTV), as well as the alpharetrovirus avian leukosis virus (ALV) are not inactivated on exposure to mildly acidic pH [21-24]. JSRV and ALV are only inactivated if first bound to their receptor, suggesting an essential two-step fusion mechanism of receptor binding followed by exposure to low pH [22, 24]. To test whether exposure to acid pH inactivates HERVK Env, we exposed purified VSV-HERVK to increasingly acidic pH for 30 minutes, neutralized the pH and then measured the residual infectivity. Treatment of purified VSV-HERVK particles at pH <6.0 reduced infectivity (Fig 4A). By contrast the infectivity of VSV was unaffected reflecting the reversibility of the conformational changes in VSV G when exposed to acid pH [25-27]. We next examined whether VSV-HERVK infection requires endocytosis beyond a need for acidic pH. For this purpose we bound virus to the cell surface and exposed cells to a brief pulse of acidic pH (Fig 4B). Infection was readily established demonstrating that endocytic uptake is not required and establishing a minimal requirement and a necessary order of virus attachment and acidic pH for HERV-K Env mediated entry

**Fig 4.**
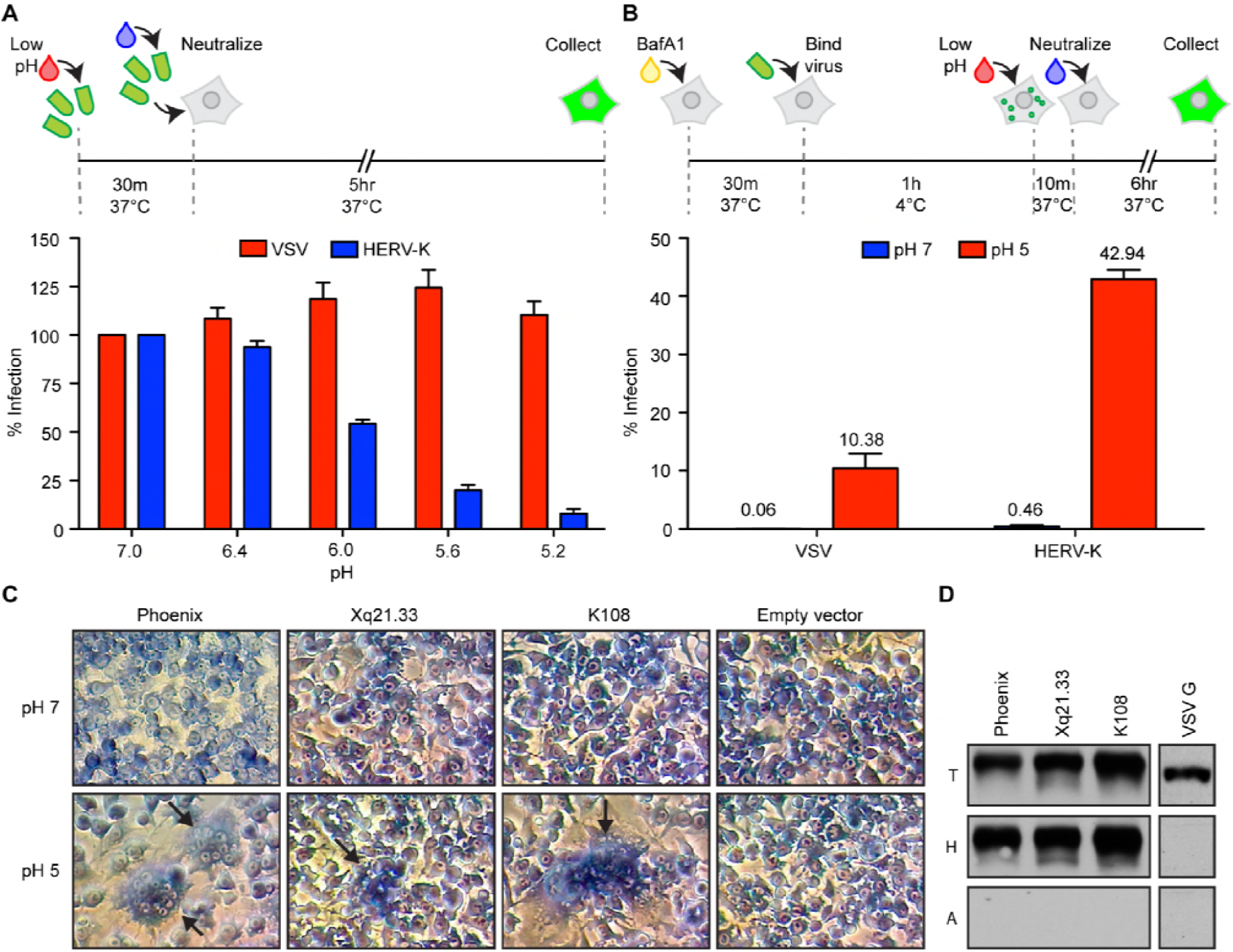
Acidic pH is sufficient to trigger HERV-K Env. (A) HERV-K Env is inactivated by exposure to acidic pH. VSV-HERVK and VSV were incubated in buffer at the indicated pH for 30 minutes at 37°C. Samples were returned to neutral pH before infecting BSRT7 cells. Cells were collected 5 hours post infection and percent GFP-expressing cells quantified by flow cytometry. Values are normalized to the pH 7 condition. Error bars represent SEM from three independent experiments. **(B)** VSV-HERVK fuses at the plasma membrane when treated with acidic pH. BSRT7 cells were pre-treated with bafilomycinA1 prior to binding virus at 4°C. Cells were treated with buffer at pH 7 or 5. Unbound virus was washed off and cells were collected 6 hours post-infection. Percent infected cells was normalized to cells not treated with bafilomycinA1. Error bars represent SEM for three independent experiments. **(C)** Endogenous HERV-K Envs are fusogenic at acidic pH. BSRT7 cells were transfected with *envs* from Phoenix, Xq21.33, and HERV-K 108 and subsequently exposed to buffer at the indicated pH. Syncytia are highlighted with arrows. Data are from a single representative experiment. **(D)** Endogenous HERV-K Envs bind heparin. 293T cells were transfected with the *envs* from Phoenix, Xq21.33, HERV-K 108, or VSV-G. Cell lysates were incubated with either heparin (H) or protein A (A) beads and bound protein analyzed by Western blot against HERV-K Env and VSV-G. T: 10% of total input. Data are from a single representative experiments.

To determine whether heparin binding and acid pH triggered fusion are retained by distinct HERVK Env sequences we compared the Phoenix Env sequence with that of two distinct genomic copies, K108 and Xq21.33 [5, 28] (S7 Fig). All three Env sequences mediate acid pH dependent cell-cell fusion, although the relative fusogenicity of Xq21.33 is reduced (Fig 4D). Using lysates of cells overexpressing the individual envelope proteins we also demonstrate that the 3 HERV K Envs, but not VSV G, are specifically bound by heparin beads (Fig 4D). These results underscore that both heparin binding and acid pH triggered fusion are shared properties of multiple HERV-K envelope sequences present in the genome.

## Discussion

The major conclusion of this study is that heparan sulfate is a direct HERV-K Env attachment factor. Binding of HERVK Env to heparan sulfate is most sensitive to the loss of 2-O sulfation, implicating this modification in attachment. Combined with earlier work we posit the following model for the entry of the retrovirus HERV-K (Fig 5). Following binding to cell surface heparan sulfate, virus is taken up into cells in a dynamin-dependent, clathrin independent manner with subsequent acidification of the endosome leading to membrane fusion and productive infection. This model is reminiscent of the sialic-acid binding and acid pH requirement for productive influenza virus entry. We cannot, however, rule out the possibility that HERV-K entry may require additional host factors not identified through the haploid genetic screening approach – such as essential host genes, or those with redundant function. The low pH-mediated inactivation of VSV-HERVK, and lack of identification of endosomal factors other than acidic pH, raise the possibility that heparan sulfate may act directly as a receptor. Regardless of whether binding to heparan sulfate is sufficient to fulfill both attachment and receptor functions, this study defines heparan sulfate as an important host factor for HERV-K Env-mediated cell entry. The ability of HERVK Env to bind heparan sulfate underscores that such binding is an ancient property of viruses.

**Fig 5.**
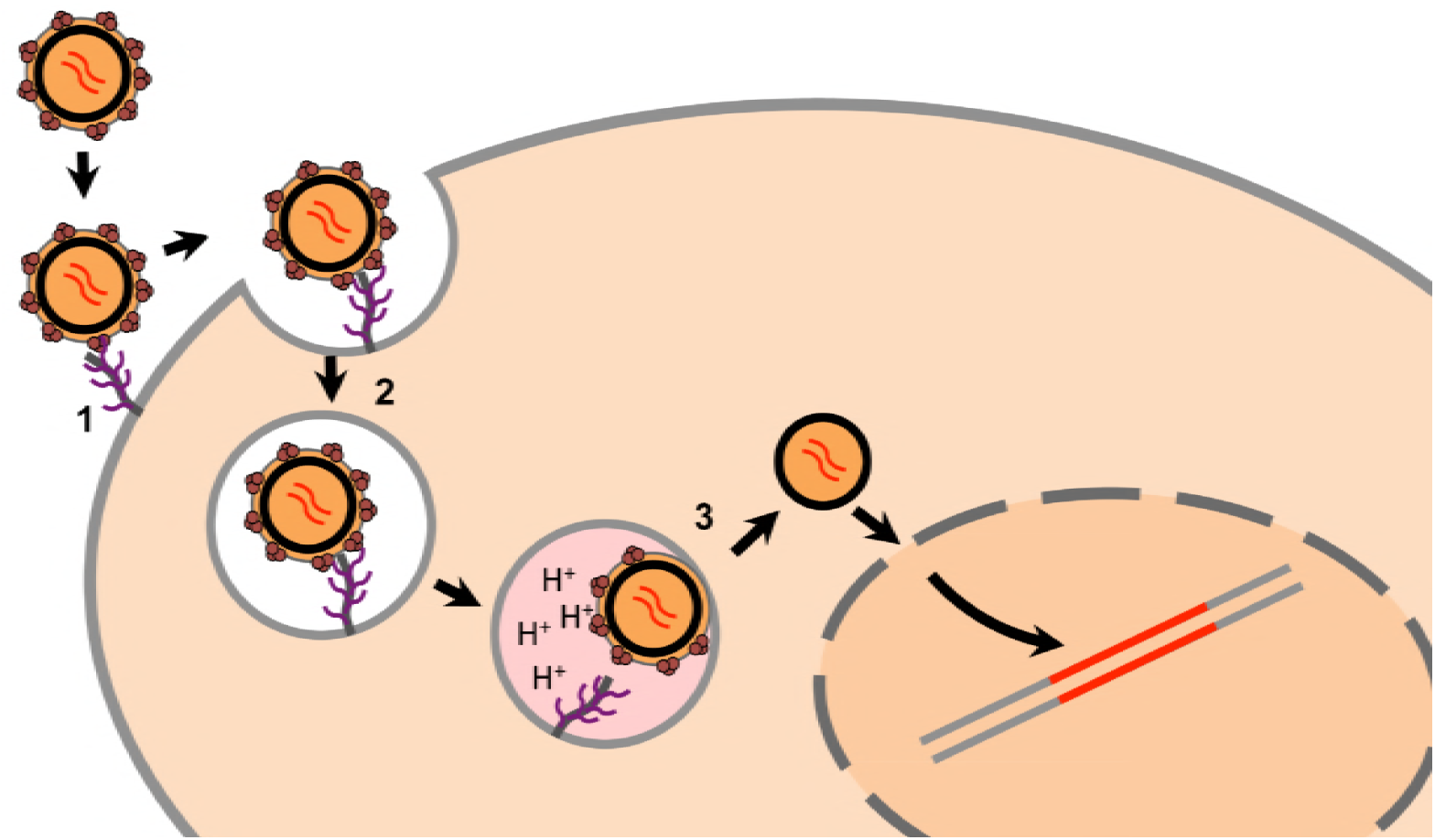
Proposed model of HERV-K entry. We propose a 3-step model for HERV-K entry: 1. HERV-K binds heparan sulfate on the cell surface to attach to the cell. 2. The virus is taken up by dynamin-dependent, clathrin-independent endocytosis. 3. Exposure to low pH following endosomal acidification triggers Env to fuse the viral and cellular membranes, releasing the viral core into the cytoplasm.

For many viruses, heparan sulfate binding reflects an adaptation to growth in cell culture. By contrast, heparan sulfate binding is an intrinsic property of the original HERV-K Env because the *env* sequences we used are not derived from viruses grown in cell culture, and such binding is apparent for multiple HERV-K Envs as they exist in the genome and at least one putative ancestral sequence. Several extant retroviruses including the prototype foamy virus (PFV), MMTV, and human T-lymphotropic virus 1 (HTLV-1) bind heparan sulfate [29-31]. MMTV requires engagement of transferrin receptor [32] and HTLV-1 requires neuropilin 1 and glucose transporter 1 [33, 34] for entry. Proteinaceous receptors for PFV have not been identified, and like HERV-K it has proved difficult to identify cell types that are refractory to entry. Although we cannot rule out the presence of an additional unidentified host factor for HERVK entry, the demonstration that acid pH alone can trigger HERVK Env suggests that engagement of such a second molecule may not be essential for infection. We do, however, observe some infectivity in cells lacking heparan sulfate by its genetic or chemical inhibition demonstrating that molecules other than heparan sulfate facilitate cell entry.

Our conclusions are based on the results from a genetic screen performed using VSV-HERVK, combined with genetic, biochemical and cell biological follow up experiments to validate the importance of heparan sulfate. In our prior work with VSV recombinants containing heterologous envelope proteins we have always validated our findings using the respective wild type virus. For HERV-K such validation experiments were not possible because the reconstructed viruses replicate poorly. Lentiviral particles pseudotyped with HERV-K Env have been described but they also produce low viral titers, ranging from approximately 60-1000 infectious units ml^-1^ as determined using a spinoculation based infectivity assay [7-9]. We obtain similar titers of pseudotyped lentiviruses without such spinoculation - 179-517 infectious units ml^-1^. Those titers are substantially below the 3×10^7^ infectious units ml^-1^ of VSV-HERVK [10], limiting the utility of the lentiviral pseudotypes in such validation experiments. Nevertheless, we carried out experiments using such lentiviral pseudotypes using Env null “bald” particles as a stringent background infectivity control. Using such lentiviral pseudotypes we observe trends similar to those with VSV-HERVK when heparan sulfate biosynthesis pathways are manipulated (S8 Fig). We therefore cannot rule out the possibility that contributions of particle geometry and glycoprotein density might influence the entry of VSV-HERVK into cells in a manner that does not fully recapitulate that of wild type HERV-K virus. We cannot know the glycoprotein density on HERV-K – it may range from the low levels observed for HIV to the high levels on MMTV [35]. Such considerations do not, however, affect the major conclusions of this study as evidenced by the fact that biochemically pure wild type HERV-K Env binds heparin, and heparin binding and acid pH triggered fusion are properties of three distinct HERV-K Env sequences.

HERV-K(HML-2) proviruses are present in all humans and HERV-K Env is expressed during a number of diseases, including viral infection, cancer, and autoimmune diseases [6, 36]. While these Envs are unlikely to be fusogenic at normal extracellular pH, they will likely act as heparan sulfate binding proteins on the cell surface and would, in principle foster contacts with the extra cellular matrix through heparan sulfate engagement. Heparan sulfate is involved in a multitude of physiological functions, from cell adhesion and migration to cell signaling [37-40]. Heparan sulfate proteoglycans have been implicated in cancer invasion and metastasis, often through dysregulation of cell signaling pathways [37]. Heparan sulfate binding by HERV-K Env could thus play a role in these processes. Overexpression of HERV-K Env on cancer cells could facilitate invasion and metastasis through binding heparan sulfate on the surface of neighboring cells or the extracellular matrix. HERV-K Env binding heparan sulfate proteoglycans may also disrupt normal signaling cascades in which these proteoglycans are involved.

As HERV-K is a relatively young group of ERVs, the ultimate fate of HERV-K Env is not yet fixed. Several HERV envelopes have been coopted throughout evolution to perform important functions for the host. These include the syncytins, which are essential for placentation [41], HERV-T Env, which has antiviral properties and may have contributed to its own extinction [42], and HEMO, a recently identified Env product that is shed in the blood of pregnant women [43]. HERV-K Env is known to be expressed in healthy tissues as well, including stem cells and during early stages of embryogenesis [36, 44]. It remains to be determined whether there is any physiological consequence of heparan sulfate binding by HERV-K Env in instances when it is actively expressed.

The conservation of heparan sulfate throughout metazoans and its ubiquitous expression presents no barrier for this ERV to enter into germ cells – a step essential for its endogenization – and implies that other steps of the HERV-K replication cycle result in the observed species tropism. The broad distribution of heparan sulfate is consistent with findings for other endogenous retroviruses. These include murine leukemia virus and MMTV, which utilize receptors that are broadly expressed in their respective host, and which exist as both endogenous and exogenous viruses [45-51]. This also holds true for other extinct primate endogenous retroviruses, chimp endogenous retrovirus 2 (CERV2) and human endogenous retrovirus T (HERV-T) [42, 52]. Perhaps the great majority of endogenous retroviruses were able to colonize the germline because their broad tropism allowed access to germ cells. Germline integration and endogenization would become chance events by such “promiscuous” viruses, rather than viruses that specifically target germ cells.

## Materials and Methods

### Cell lines, viruses, and plasmids

BSRT7 cells (a kind gift from U.J. Buchholz[53]) and 293T cells (ATCC CRL-3216; American Type Culture Collection, Manassas, VA) were grown in Dulbecco’s modified Eagle’s medium (DMEM) supplemented with 10% fetal bovine serum (FBS) and maintained at 37°C and 5% CO_2_. HAP1 cells (a kind gift from Thijn Brummelkamp [12]) were grown in Iscove’s modified Dulbecco’s medium (IMDM) supplemented with 10% FBS and maintained at 37°C and 5% CO_2_. All cell lines were tested to be free of mycoplasma. VSV-HERVK^+^, referred to throughout this manuscript as VSV-HERVK, and VSV-eGFP were generated as previously described [10, 54]. VSV-HERVK encodes the env from the Phoenix consensus sequence[8], where the cytoplasmic tail has been replaced with that of VSV G. Both viruses express eGFP from the first position in the genome. Viruses were grown on BSRT7 cells. Viruses were tittered by plaque assay on BSRT7 cells and flow cytometry on HAP1 cells.

pCAGGS-PhoenixEnv and pGEM-PhoenixEnv were previously generated [10]. We synthesized codon-optimized versions of HERV-K 108 *env* and Xq21.33 *env* (Genscript, Piscataway, NJ) and subsequently cloned them into pCAGGS and pGEM to generate pCAGGS-HERVK108Env, pCAGGS-Xq21.33Env, pGEM-HERVK108Env, and pGEM-Xq21.33Env, respectively. Plasmids containing the cDNA for *EXT1* and *SLC35B2* were obtained from the Dana-Farber plasmID repository (Dana-Farber/Harvard Cancer Center DNA Resource Core, Boston, MA). These cDNAs were amplified with primers to add a C-terminal HA tag and were cloned into pQCXIN (Clontech, Mountain View, CA). lentiCas9-Blast [55] (Addgene plasmid #52962), lentiGuide-Puro [55] (Addgene plasmid #52963), lentiCRISPRv2 [55] (Addgene plasmid #52961) and pX330 [56] (Addgene plasmid #42230) were a gift from Feng Zhang.

### Haploid genetic screen with VSV-HERVK

HAP1 cells were mutagenized with a genetrap retrovirus as described [13]. Approximately 10^8^ cells were infected with VSV-HERVK at a multiplicity of infection (MOI) of 3 infectious units (IU) per cell. Infection was allowed to proceed for several days, after which a second round of infection was performed to kill remaining susceptible cells. Genomic DNA was isolated from surviving cells and used to prepare a library for Illumina deep sequencing and reads analyzed as described [13]. Inactivating insertion sites (mapping to exons or in sense orientations in introns) in the VSV-HERVK selected cells (180,655 unique insertions) were compared to that of a control data set from unselected cells (2,161,301 unique insertions). *P* values for enrichment in the selected set versus control set were calculated using a Fisher’s exact test. Significance scores are reported as the inverse log of the *p* value. Genes with insertions were also analyzed to identify bias in the direction of insertion of the genetrap sequence within introns. Insertions in the forward direction are inactivating.

### Generation of knockout cells

To generate *CREBBP*^KO^ cells, a guide RNA targeting the Histone Acetyl Transferase domain of CREBBP (5’-GGAGGTTTTTGTCCGAGTGG-3’) was cloned into pX330 creating pX330-*CREBBP*. HAP1 cells were co-transfected with pX330-*CREBBP* and a plasmid containing an expression cassette for a guide RNA targeting the zebrafish *TIA* gene (5’-GGTATGTCGGGAACCTCTCC-3’) followed by a CMV promotor sequence driving expression of a blasticidin resistance gene flanked by two *TIA* target sites [57]. Co-transfection of these plasmids resulted in the incorporation of the blasticidin resistance cassette at the site of the targeted *CREBBP* locus. Four days after DNA transfection, the culture medium was supplemented with blasticidin (30 µg/mL). A single cell, blasticidin-resistant clone (C4C2) was expanded and disruption of *CREBBP* was verified by Sanger sequencing and Western blot for protein expression (using anti-CREBBP; clone C-20, Santa Cruz Biotechnology, Santa Cruz, CA).

To generate *MYO10*^*KO*^ and *SORT1*^*KO*^ cells, we first generated HAP1 cells stably expressing *Streptococcus pyogenes* Cas9 (HAP1-Cas9). Lentivirus was generated by transfecting 293T cells with lentiCas9-Blast, pCD/NL-BH*DDD [58] and pCAGGS-VSVG. HAP1 cells were transduced with the lentivirus and selected with 5μg/ml blasticidin. Guide RNAs targeting exon 20 of *MYO10* (5’-CGCCTGGTCAAACTGGTC-3’) and exon 3 of *SORT1* (5’-CAGTCCAAGCTATATCGA-3’) were individually cloned into lentiGuide-Puro. Lentivirus was generated by transfecting 293T cells with lentiGuide-MYO10sgRNA or lentiGuide-SORT1sgRNA, pCD/NL-BH*DDD and pCAGGS-VSVG. HAP1 cells were transduced with the lentivirus and selected with 2 μg/ml puromycin. Single cell clones were screened by Western blot for expression of myosin-X and sortilin using anti-MYO10 (HPA024223; Sigma-Aldrich, St. Louis, MO) and anti-sortilin (ab16640; Abcam, Cambridge, MA) antibodies. To generate *EXT1*^KO^ cells, guide RNAs targeting two sequences in exon 1 of *EXT1* (5’-GGCCAGAAATGATCCGGACT-3’ and 5’-GCACAACGTCCTCCCCGTTA-3’) were individually cloned into pX330. The plasmids were cotransfected along with pLPCX-Puro (Clontech) into HAP1 cells using lipofectamine 3000 transfection reagent (Life Technologies, Grand Island, NY). Cells were selected with 2 μg/ml puromycin. Genomic DNA from single cell clones was isolated using DirectPCR lysis reagent (Viagen Biotech, Los Angeles, CA) and screened by PCR for an approximately 700 base pair deletion using the following primers: 5’-GAGTTGAAGTTGCCTTCCCG-3’ and 5’-AGGCTTTTCAGTTTGCCCGA-3’. To generate *SLC35B2*^KO^cells, a guide RNA sequence targeting exon 4 of *SLC35B2* (5’-GCTTTCCCATCAGCATGACA-3’) was cloned into the lentiCRISPRv2 plasmid [55]. Lentivirus was generated by transfecting 293T cells with lentiCRISPRv2-SLC35B2sgRNA, pCD/NL-BH*DDD and pCAGGS-VSVG. HAP1 cells were transduced with the lentivirus and selected with 2 μg/ml puromycin. Single cell clones were screened by flow cytometry for reactivity with 10E4 (370255-1, AMS biotechnology, Cambridge, MA), an antibody specific for sulfated heparan sulfate. After selection and single cell cloning, all cells were maintained in IMDM+10% FBS without puromycin.

*B4GALT7*^KO^ HEK293T cells were generated by CRISPR/Cas9-mediated genome editing essentially as described by Langereis et al [59] with two guide RNAs targeting B4GALT7’s active site encoded in exon 5 (gRNA 1: 5’-ATGGGATGTCCAACCGCTTC-3’ and 5’-GAGTTCTACCGGCGCATTAA-3’). Single cell clones were isolated by fluorescence activated cell sorting (FACS) and genotyped by PCR and DNA sequencing. Loss of heparan sulfate expression was confirmed by flow cytometry with antibody 10E4.

### Reconstitution of knockout cells with EXT1 or SLC35B2

Murine leukemia virus (MLV) carrying either EXT1-HA or SLC35B2-HA was generated by transfecting 293T cells with pCS2-MGP (Moloney MLV gag/pol expression vector [60]), pCAGGS-VSVG, and either pQCXIN-EXT1HA, pQCXIN-SLC35B2HA or pQCXIN empty vector. *EXT1*^KO^ cells were transduced with MLV-EXT1HA or MLV-empty vector to generate *EXT1*^KO^+EXT1-HA or *EXT1*^KO^ Neo^r^ control, respectively. *SLC35B2*^KO^ cells were transduced with MLV-SLC35B2HA or MLV-Empty vector to generate *SLC35B2*^KO^+SLC35B2-HA or *SLC35B2*^KO^ Neo^r^ control, respectively. Cells were selected with 1.5 mg/ml G418. *SLC35B2*^KO^+SLC35B2-HA cells were further subcloned and single cell clones were screened for heparan sulfate expression using the 10E4 antibody. All reconstituted cells were maintained in IMDM with 10% FBS and 1.5 mg/ml G418.

### Heparan sulfate staining

Cells were collected with 10mM EDTA in PBS and subsequently resuspended in PBS with 1% BSA and BD human Fc block (1:25 dilution; 564220, BD Biosciences, San Jose, CA). Cells were incubated with either the 10E4 monoclonal antibody to heparan sulfate (1:400), or mouse IgM (1:200; ab18401, Abcam, Cambridge, MA) for 1 hour (h) on ice. Cells were washed twice with PBS+1%BSA and incubated with goat anti-mouse IgG/IgM alexafluor 488 (1:500; A10680, Molecular Probes, Eugene, OR) for 1h on ice. Cells were washed twice and fixed in 2% paraformaldehyde (PFA). Fluorescence was measured using a FACSCalibur instrument (Cytek Development, Freemont, CA) and analyzed using FlowJo software (FlowJo Inc, Ashland, OR).

### Infectivity experiments

WT HAP1, *MYO10*^*KO*^, *CREBBP*^*KO*^, *SORT1*^KO^, *EXT1*^*KO*^, *EXT1*^*KO*^ Neo^r^ control, *EXT1*^*KO*^+EXT1-HA, *SLC35B2*^*KO*^, *SLC35B2*^*KO*^ Neo^r^ control, or *SLC35B2*^*KO*^+SLC35B2-HA cells were infected with VSV-HERVK or VSV-eGFP at an MOI of 1 IU/cell. Cells were collected 5h post-infection (pi) and fixed in 2% PFA. eGFP fluorescence was measured using a FACSCalibur instrument and the percentage of eGFP-positive cells, as well as mean fluorescence intensity was quantified using FlowJo software. The number of individual clones tested for each cell line are as follows: *MYO10*^KO^ (n=2), *CREBBP*^KO^ (n=1), *SORT1*^KO^ (n=3), *EXT1*^KO^ (n=3), *SLC35B2*^KO^ (n=1). Data are shown from multiple independent replicates in a single clonal cell line for each gene. Data are represented as fold difference in percent eGFP-positive cells or MFI compared to WT cells and normalized to VSV infectivity in each cell type. Error bars represent standard error of the mean from at least three independent biological replicates.

### Sodium chlorate treatment and infectivity

BSRT7 cells were passaged in sulfate-free Joklik modified minimal essential medium (M8028, Sigma-Aldrich, St. Louis, MO) with 10% FBS with or without 50mM sodium chlorate for at least two passages prior to seeding for infection. Cells were infected with either VSV-HERVK or VSV at an MOI of 1 particle forming unit (PFU)/cell. Cells were collected 5hpi and fixed in 2% PFA and fluorescence measured using a FACSCalibur instrument. The % eGFP-positive cells and MFI were quantified using FlowJo software. Data are represented as fold difference in % eGFP-positive cells or MFI normalized to VSV infected cells. Error bars represent standard error of the mean from three independent biological replicates.

### Inhibition of infection by soluble glycosaminoglycans

Heparin (H3393, Sigma-Aldrich), heparan sulfate (AMS GAGHS01, AMS bioscience), chondroitin sulfate A (C9819, Sigma-Aldrich) and dermatan sulfate (C3788, Sigma-Aldrich) were reconstituted in PBS. 1 μg of purified VSV-HERVK or VSV was incubated with the compounds at the indicated concentration in PBS for 1h at 37°C and then added to BSRT7 cells. Cells were incubated with virus and compound for 1h at 37°C, washed with DMEM and incubated for 4h at 37°C. Cells were collected, fixed with 2% PFA and fluorescence measured using a FACSCalibur instrument. The % eGFP-positive cells were quantified using FlowJo software and normalized to infectivity with no compound. Error bars represent standard error of the mean from three independent biological replicates.

### Imaging of HERV-K attachment to cells

Gradient purified VSV-HERVK particles were labeled with AlexaFluor 647 and VSV particles were labeled with AlexaFluor 594, as described [61]. WT HAP1 or *SLC35B2*^*KO*^ cells were pre-stained with calcein (diluted 1:1000; C3099, Molecular Probes, Eugene, OR) and NucBlue live cell stain (1:50, C34552, Molecular Probes) in IMDM for 30 minutes (min) at 37°C, followed by blocking in 1% BSA in IMDM for 30 min at 37°C. Labeled VSV-HERVK and VSV were added together to the cells. Cells were incubated with virus at either 37°C for 15 min or at 4°C for 1 h. Samples were fixed in 2% PFA and mounted with ProLong Gold (P10144, Molecular Probes). Samples were imaged using a Marianas system (Intelligent Imaging Innovations, Denver, CO) based on a Zeiss observer microscope (Carl Zeiss Microimaging, Thornwood, NY) outfitted with a 64 CSU-22 spinning-disk confocal unit (Yokogawa Electric Corporation, Musashino, Tokyo, Japan) and a 63x (Plan-Apochromat, NA 1.4; Carl Zeiss Microimaging) objective lens. Excitation wavelengths were 561 nm for AF594 and 660 nm for AF647. SlideBook 6.0 (Intelligent Imaging Innovations) was used to command the hardware devices and visualize and export the acquired data. Subsequent image analysis was conducted using ImageJ (National Institutes of Health). Briefly, cellular cytoplasmic areas were approximated by manually tracing the 2D cellular outline based on the calcein staining and determining its area. To simplify visualization, calcein aggregates were eliminated using the Remove Outliers tool in ImageJ. Bound VSV and VSV-HERVK particles were counted for each cell, excluding large aggregates. Particle binding per area unit was calculated by dividing particle counts by the calculated areas. Data are represented as box plots indicating the median values, first and third quartiles, minima and maxima. Outliers were defined as those points 1.5 times the interquartile range, and severe outliers as those 3 times the interquartile range. Data are from multiple images from a single experiment. N values are as follows: for the 37°C experiment, 29 WT cells were counted, with 189 VSV particles and 492 VSV-HERVK particles; 62 *SLC35B2*^KO^ cells were counted with 252 VSV particles and 220 VSV-HERVK particles. For the 4°C experiment, 43 WT cells were counted, with 629 VSV particles and 793 VSV-HERVK particles; 72 *SLC35B2*^KO^ cells were counted with 790 VSV particles and 377 VSV-HERVK particles.

### Virus and cell lysate heparin pull-downs

For all pull-downs, heparin beads (H6508, Sigma-Aldrich) and protein A beads (P2545, Sigma-Aldrich) were washed 3 times in the corresponding buffer and blocked for 1 h in PBS with 1% BSA. BSRT7 cells were transfected with pCAGGS-PhoenixEnv, pCAGGS-HERVK108Env, or pCAGGS-Xq21.33Env. At 24h post-transfection, cells were lysed in TNE buffer (50mM Tris, 2mM EDTA, 150mM NaCl,) supplemented with 1% tritonX-100 and 200 μl lysate was incubated with 50 μl of either heparin or control protein A beads at 4°C for 2 h. Beads were washed 5 times in TNE+1% tritonX-100 and bound proteins eluted by boiling in 2X SDS loading buffer for 5 min. Samples were loaded on a 10% (wt/vol) polyacrylamide gel and transferred to nitrocellulose membrane which were blocked with 5% milk in PBS+0.1% Tween-20 and subsequently blotted with anti-HERV-K Env antibody (1:2000; HERM-1811-5, Austral biological, San Ramon, CA) followed by goat anti-mouse horseradish peroxidase (HRP) antibody (1:5000; Sigma-Aldrich). Membranes were incubated with ECL reagent (Thermo-Fisher Scientific, Waltham, MA) and signal was detected by film (Denville Scientific, Holliston, MA). Data shown are from a single representative experiment from 3 biological replicates.

For virus pull-down we incubated 10 μg of purified virus +/-50 μg/ml heparin in TNE buffer with 1% BSA. Complexes were collected by incubation with 50 μl heparin or protein A beads (4°C rotating for 2h), and washed 5 times in TNE buffer. Bound virions were eluted and analyzed by Western blot as described above. Membranes were blotted with antibodies against HERV-K Env, VSVG (1:10,000; V5507, Sigma-Aldrich), or VSVM (1:5000; 23H12, a kind gift from Doug Lyles[62]) followed by goat anti-mouse HRP antibody (1:5000; Sigma-Aldrich). Membranes were incubated with ECL reagent and signal was detected by film. Data shown are from a single representative experiment from 3 biological replicates.

### Generation and characterization of HERV-K Env truncations

N-terminal truncations (N1-N7) were designed as outlined in S5 Fig to determine the appropriate boundary between the signal peptide and SU domain. DNA fragments containing the truncated versions of the envelope were cloned into a modified pVRC8400 expression vector, which uses tissue plasminogen activator signal sequence [63]. These *env* sequences with the new signal peptide were sub-cloned into pGEM3, under the control of a T7 polymerase promoter. Env truncations were screened for expression, proteolytic processing, fusogenicity, and pH dependency of fusion. Each screening experiment was a single replicate. Western blot analysis and cell-cell fusion experiments were performed as described [10]. Briefly, BSRT7 cells were infected with VVT7.3 [64], a vaccinia virus encoding the T7 RNA polymerase as source of transcriptase. The cells were subsequently transfected with the HERV-K *env* expression plasmids or an empty vector control. At 18h post-transfection, cells were either harvested for Western blot analysis against HERV-K Env TM subunit (Austral biological) and Actin (Abcam), or treated with phosphate-buffered saline (PBS) at the indicated pH for 20 min at 37°C, at which point the cells were washed and standard growth medium was added. The cells were incubated for 4 h at 37°C and subsequently imaged. Truncations N4 and N5 had similar expression, processing, fusogenicity, and pH-dependency as WT.

C-terminal truncations based on N4 and N5 were tagged with a C-terminal HA tag, 3C protease cleavage site, and a tandem His_8X_-His_6X_ tag and cloned into pVRC8400. Recombinant protein was produced by transient transfection of 293T cells using Lipofectamine 2000 (Life Technologies), per manufacturer’s protocol. Three days post-transfection supernatants were harvested and clarified from cellular debris by low-speed centrifugation. HERV-K SU was purified by passage over Co-NTA agarose (Clontech) and concentrated with an Amicon Ultra-4 filter (Millipore, Billerica, MA). Purified protein was run on both reducing and non-reducing SDS-PAGE (4-20% polyacrylamide gel, 4561096; Bio-rad, Hercules, CA) followed by Coomassie staining. Proteins were screened for the following criteria: 1. Expression: If a band of appropriate size was observed on a reducing gel. 2. Solubility: the absence of major aggregate bands under non-reducing conditions. 3. Monomeric: only proteins without evidence of major aggregation were subject to size exclusion chromatography. Proteins for which a discrete peak in the A280 trace corresponding to the approximate size of monomeric SU and produced a single band of the appropriate size on a non-reducing SDS-PAGE were deemed to produce monomeric species.

### Production of soluble HERV-K SU

A truncated version of the codon optimized HERV-K Phoenix SU domain, encoding residues 96-433 (residue 1 being the initiating methionine) was synthesized by Integrated DNA Technologies, Inc. to included a C-terminal HA tag, 3C protease cleavage site, and a tandem His_8X_-His_6X_ tag. This cDNA was cloned into the modified pVRC8400 expression vector [63]. Recombinant protein was produced by transient transfection of 293T cells using Lipofectamine 2000 (Life Technologies), per manufacturer’s protocol. At 3 days post-transfection supernatants were harvested and clarified from cellular debris by low-speed centrifugation and HERV-K SU purified by passage over Co-NTA agarose (Clontech) followed by gel filtration chromatography on Superdex 200 (General Electric Healthcare, Piscataway, NJ) in 10 mM Tris-HCl, 150 mM NaCl at pH 7.5. Three major peaks were observed: an aggregate of SU that eluted in the void volume of the column, a dimeric species that could be reduced into monomers by addition of a reducing agent (likely the result of a disulfide bond linking two monomers), and a major peak containing a homogeneous monomeric species. For binding assays, only gel filtration chromatography fractions containing the monomeric species were used.

### Purification of recombinant HA

The hemagglutinin (HA) gene of Influenza A virus A/Leningrad/360/1986(H3N2) HA (Accession number CY121277) was synthesized as a gBlock (Integrated DNA Technologies, Inc., Coralville, IA) and used as a template to amplify the globular head of HA, residues 37-319 (Hong Kong 1968 H3N2 numbering). The resulting PCR product was cloned and expressed from a baculovirus recombinant as previously described [63]. The HA head was purified by passage over Co-NTA agarose (Clontech) followed by gel filtration chromatography on Superdex 200 (GE Healthcare) in 10 mM Tris-HCl, 150 mM NaCl at pH 7.5.

### Heparin pull downs with purified SU

Briefly, 3 μg of HERV-K SU or HA was pre-incubated with 50 μg/ml heparin, heparan sulfate, condriotin sulfate A, dermatan sulfate, 2-O-desulfated heparin (AMSDSH001-2; AMS Bioscience), 6-O-desulfated heparin (AMSDSH002-6; AMS Bioscience), or no compound in 10 mMTris-HCl, 150 mM NaCl, 0.2% TritonX-100 for 1 h at 4°C, and subsequently mixed with 50 ul of heparin, protein A or cobalt beads prior to incubation for 2 h at 4°C. Beads were washed 5 times, bound proteins eluted as above, separated on a 4-20% acrylamide gel (Bio-rad) and transferred to nitrocellulose membranes. Membranes were blotted with an antibody against the HA tag (1:5000; Abcam) followed by anti-rabbit HRP antibody (1:5000; Sigma-Aldrich). Data shown are representative of 3 (Fig 3C) or 2 (Fig 3D) independent biological replicates.

### Low pH inactivation of virions

Virus was incubated in buffer (10mM Na_2_HPO_4_, 10mM HEPES, 10mM MES) at various pH (7.0, 6.4, 6.0, 5.6, and 5.2) for 30 min at 37°C. pH was neutralized by adding an excess of DMEM+10% FBS and residual viral infectivity determined by infection of BSRT7 cells. Cells were collected 5 hpi, fixed in 2% PFA and eGFP fluorescence measured using a FACSCalibur instrument. The % of eGFP-positive cells was quantified using FlowJo software and normalized to pH7 treatment controls. Error bars represent standard error of the mean from 3 independent biological replicates.

### Acid bypass of endocytosis

BSRT7 cells were treated with 100nM bafilomycin A1 (Sigma-Aldrich; B1793) for 30 min at 37°C and VSV or VSV-HERVK subsequently bound by incubating cells with virus an MOI of 5 PFU/cell. for 1h at 4°C. Bound virus was then pulsed with buffer (10mM Na_2_HPO_4_, 10mM HEPES, 10mM MES) at either pH 7 or pH 5 for 10 min at 37°C, cells were washed twice and then incubated with DMEM (+/-100nM bafilomycin A1). At 6 hpi cells were collected, fixed in 2% PFA and eGFP fluorescence measured as above. The % eGFP-positive cells in Bafilomycin treated cells is expressed relative to untreated cells. Error bars represent standard error of the mean from three independent biological replicates.

### Cell-cell fusion experiments

Cell-cell fusion experiments were performed as previously described [10]. Briefly, BSRT7 cells were infected with VTF7-3 [64], transfected with pGEM plasmids encoding the *env* of Phoenix, Xq21.33, or HERV-K 108 or empty vector and treated with a 20 min pulse of DMEM of varying pH at 18h post-transfection. Cells were washed, and incubated for 4h at 37°C in DMEM. Cells were fixed in cold methanol prior to Giemsa staining according to manufacturer’s protocol (Sigma-Aldrich). Data shown are from a single representative experiment from three biological replicates.

### Lentiviral pseudotypes infections

Lentiviruses pseudotypes were generated by transfecting 293T cells with pCD/NL-BH*DDD, pNL-EGFP/CMV-WPREDU3 [65], and either pCAGGS-PhoenixEnv, pCAGGS-VSVG, or pCAGGS empty vector (to generate bald particles). Supernatant was collected 48 hours post transfection and particle concentration was determined using a p24 (HIV-1) antigen capture kit from Advanced Bioscience Labs (Rockville, MD). Supernatant volumes were equilibrated to equal particle amounts, based on p24 values, of each pseudotype virus were used to infect 293T, 293T-*B4GALT7*^KO^, CRFK and CRFK cells treated with 50mM NaClO_3_ for two passages prior to infection. Supernatant was removed from cells 24 hours post infection and cells were collected 48 hours post infection. For NaClO_3_ treated cells, NaClO_3_ was present during the infection and subsequent incubation. eGFP fluorescence was measured using a FACSCalibur instrument and the percentage of eGFP-positive cells, as well as mean fluorescence intensity was quantified using FlowJo software. Error bars represent standard error of the mean from three independent biological replicates.

## Acknowledgments

We thank T.R. Brummelkamp for providing critical reagents and facilities as well as critical comments on this manuscript.

---

**Fig S1.**
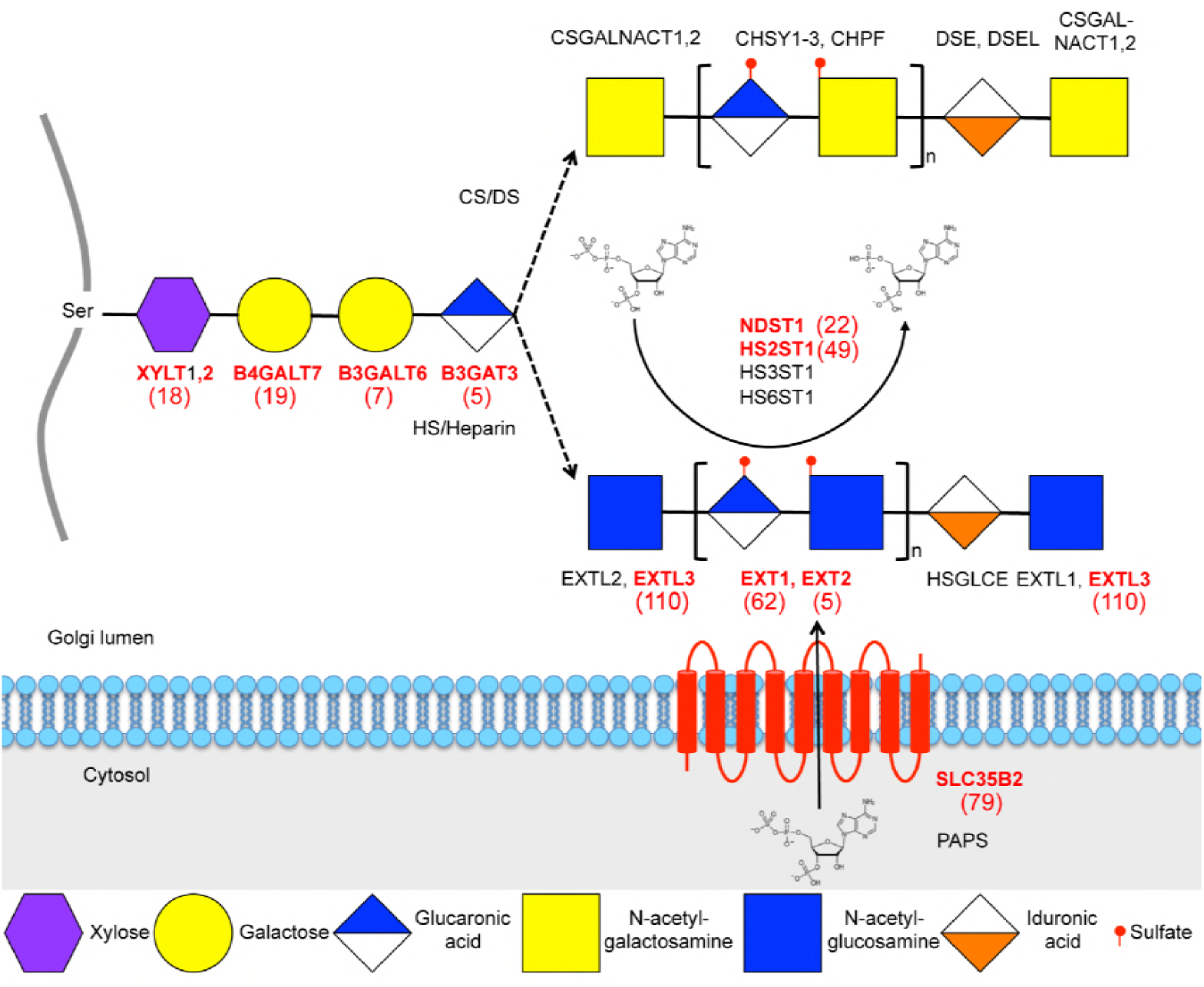
Cartoon schematic of the glycosaminoglycan (GAG) synthesis pathway. GAGs are added to a core protein (in grey). There is a core linkage of 4 sugars. The pathway then splits into the heparan sulfate/heparin pathway and the chondroitin sulfate/dermatan sulfate pathway. The enzymes that catalyze the sugar addition are written above/below the sugars. Sulfation is catalyzed by enzymes NDST1, HS2ST1, HS3ST1, and HS6ST1-3. Each enzyme adds a sulfate to a different position on the sugar. The sulfate donor, PAPS is transported into the Golgi by SLC35B2. Genes highlighted in red were identified as hits in the haploid screen. The significance score for each hit, rounded to the nearest integer, is indicated in parentheses.

**Fig S2.**
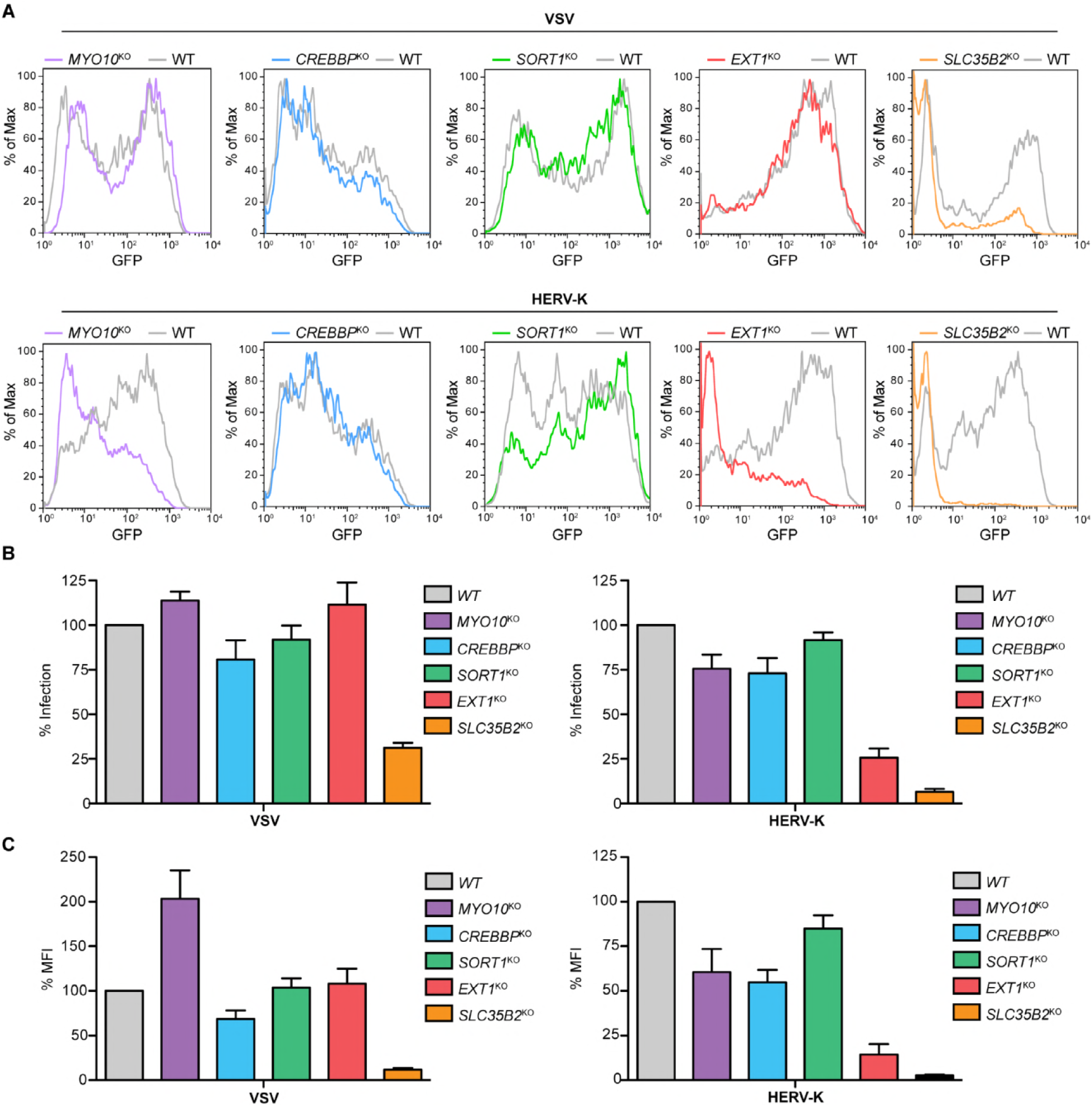
Histograms of GFP expression and infectivity in cells from Fig 1. (A) Representative histograms from experiments in Fig 1D. Top: VSV infected cells. Bottom: VSV-HERVK infected cells. Histograms are from a single representative experiment. **(B)** Infectivity of VSV and VSV-HERVK in gene edited cells. Data are from the same experiment as Fig 1D and are normalized to infectivity in WT cells. **(C)** MFI of cells in (B). MFI of all cells for each condition are normalized to that of WT cells.

**Fig S3.**
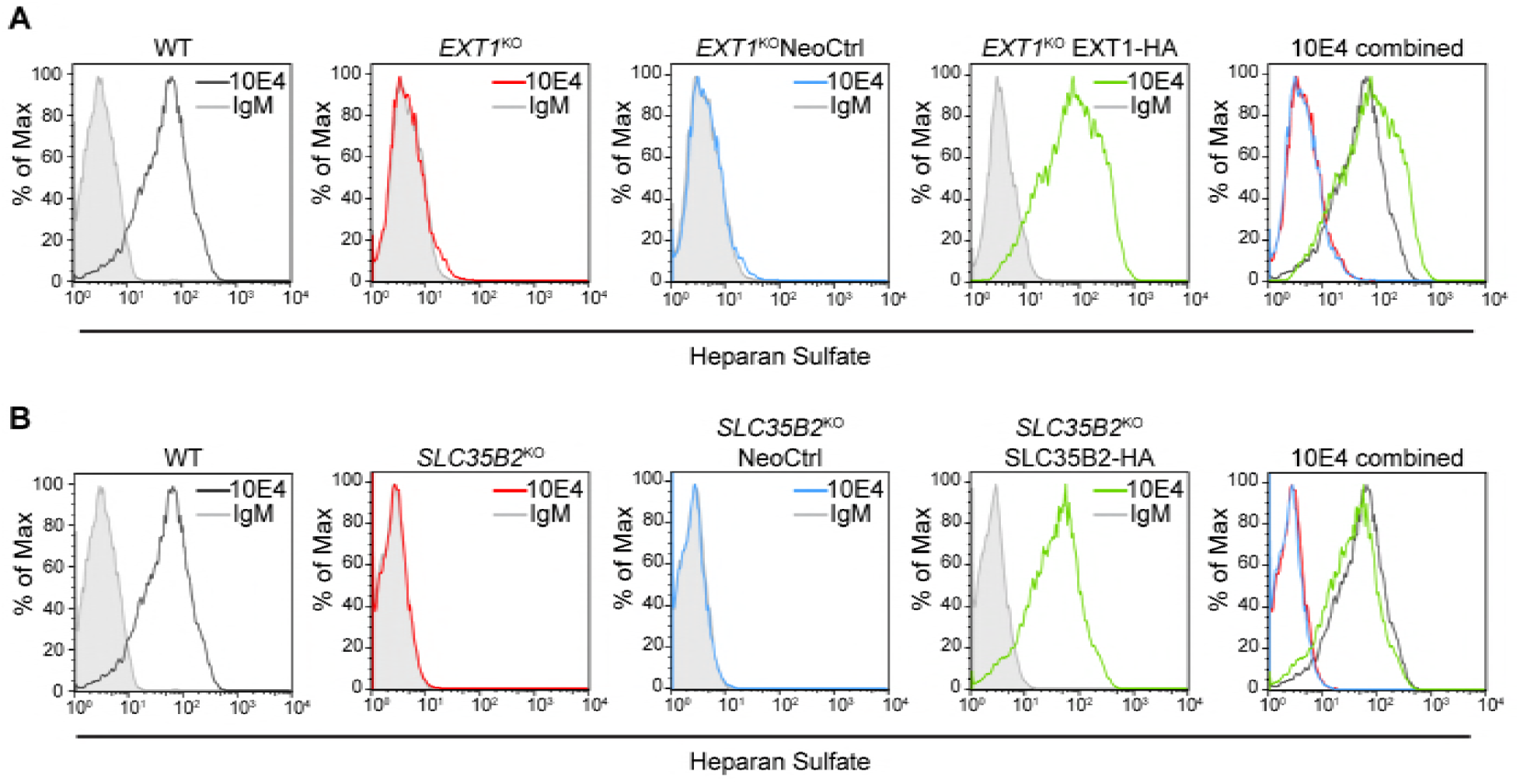
Heparan sulfate expression of. ***EXT1*^KO^ and *SLC35B2*^KO^ HAP1 cells.** The indicated cell lines were stained with 10E4, a heparan sulfate-specific antibody, or mouse IgM isotype control antibody, and analyzed by flow cytometry. **(A)** WT HAP1, *EXT1*^*KO*^, *EXT1*^*KO*^Neo^r^ Control, and EXT1^KO^+EXT1-HA cells. **(B)** WT HAP1, *SLC35B2*^*KO*^, *SLC35B2*^*KO*^Neo^r^ Control, and *SLC35B2*^*KO*^+SLC35B2-HA cells. Representative histograms are shown from a single experiment. The WT histograms in (A) and (B) are the same sample and therefore identical. Histograms are from a single representative experiment.

**Fig S4.**
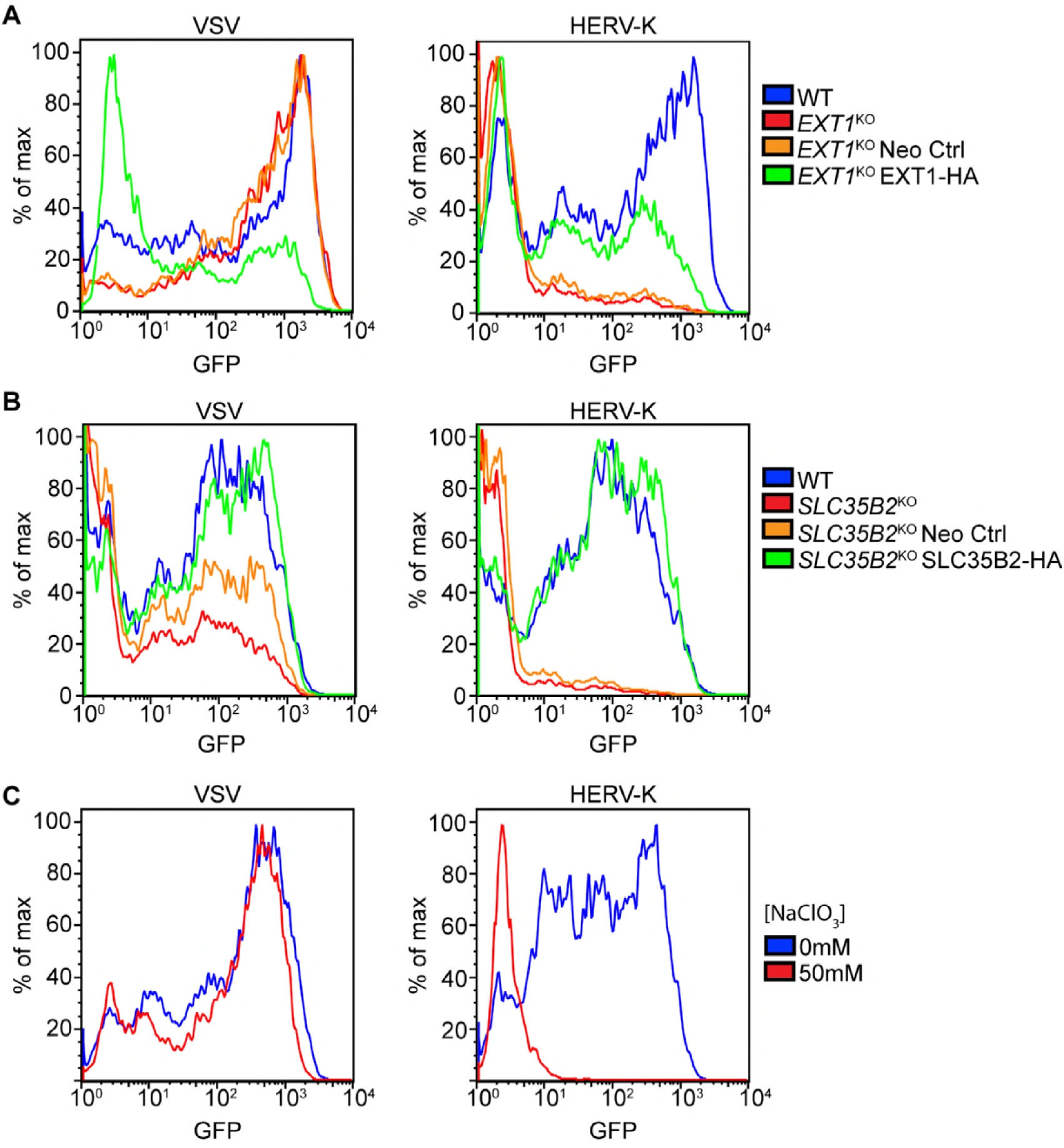
Histograms of GFP expression in cells from Fig 2. Representative histograms are shown from experiments from experiments in Fig 2A, 2B, and 2C. **(A)** *EXT1*^KO^ cells. **(B)** *SLC35B2*^KO^ cells. **(C)** Sodium chlorate treated cells. Representative histograms are shown from a single experiment.

**Fig S5.**
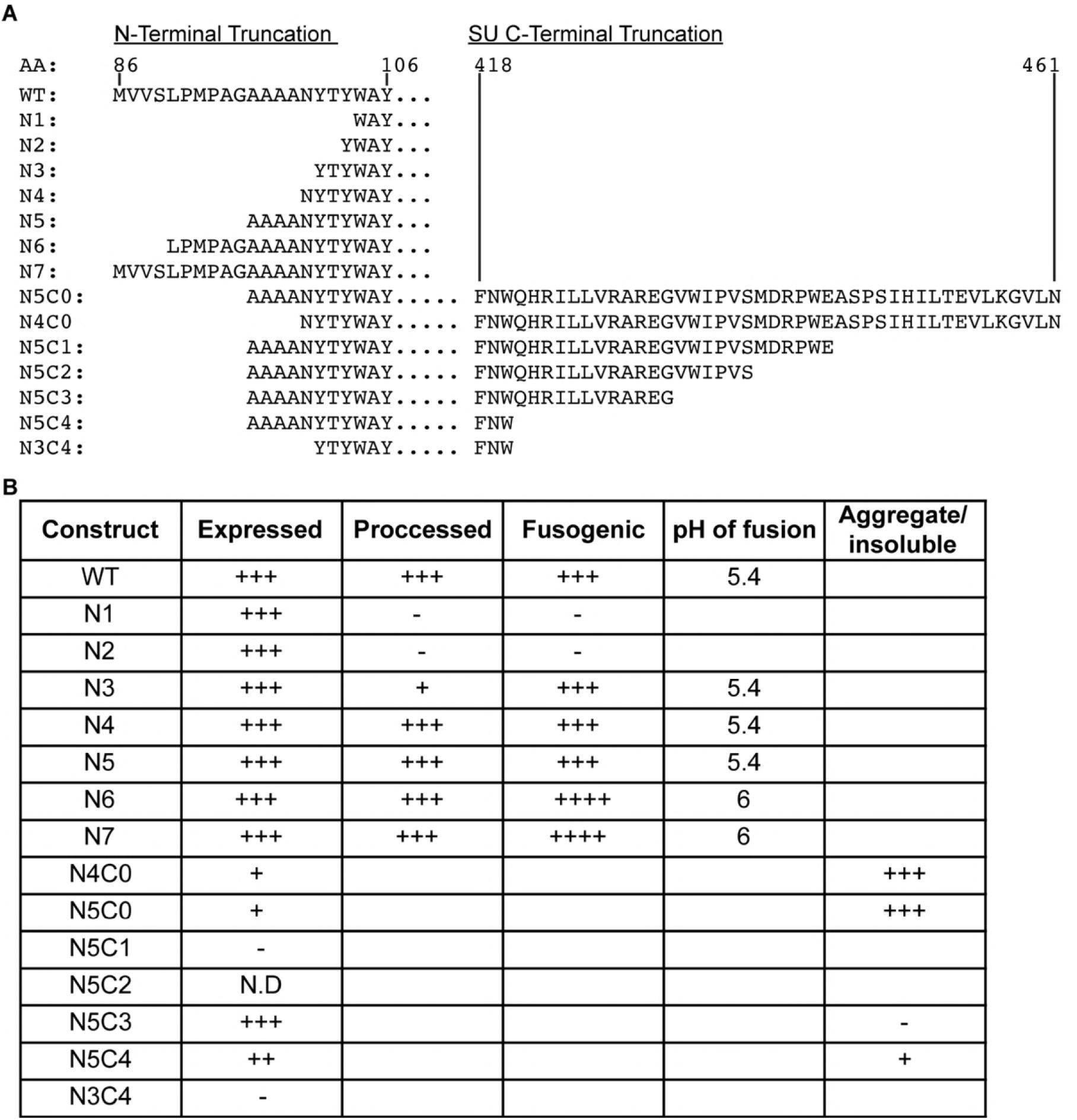
Generation and characterization of soluble HERV-K SU. (A) Schematic of the sequences of the various N-and C-terminal truncations tested. The tissue plasminogen activator signal peptide was introduced at the N-terminus of all of the truncations. N1-N7 were made in otherwise full-length sequences. C-terminal truncations were further modified with an HA-tag, a 3C protease cleavage site, and a tandem His_8X_-His_6X_ tag. Amino acid residue numbers are indicated above the sequences, with 1 being the initiating methionine. **(B)** Characteristics of HERV-K Env truncations. N1-N7 were expressed in BSRT7 cells and tested by Western blot for expression and proteolytic processing, and by cell-cell fusion assay for fusogenicity and pH dependence. C-terminal truncations were expressed in 293T cells. Protein in supernatant was isolated over cobalt resin and tested for expression, solubility and oligomerization state. N.D.: Not determined. Empty boxes: Assay not applicable to given construct. +: 1-30% of WT levels. ++: 31-60% of WT levels. +++ 61-100% of WT levels. ++++: 101-130% of WT levels. For C-terminal truncations, values are compared to N5C3. pH of fusion: Highest pH at which cell-cell fusion was observed.

**Fig S6.**
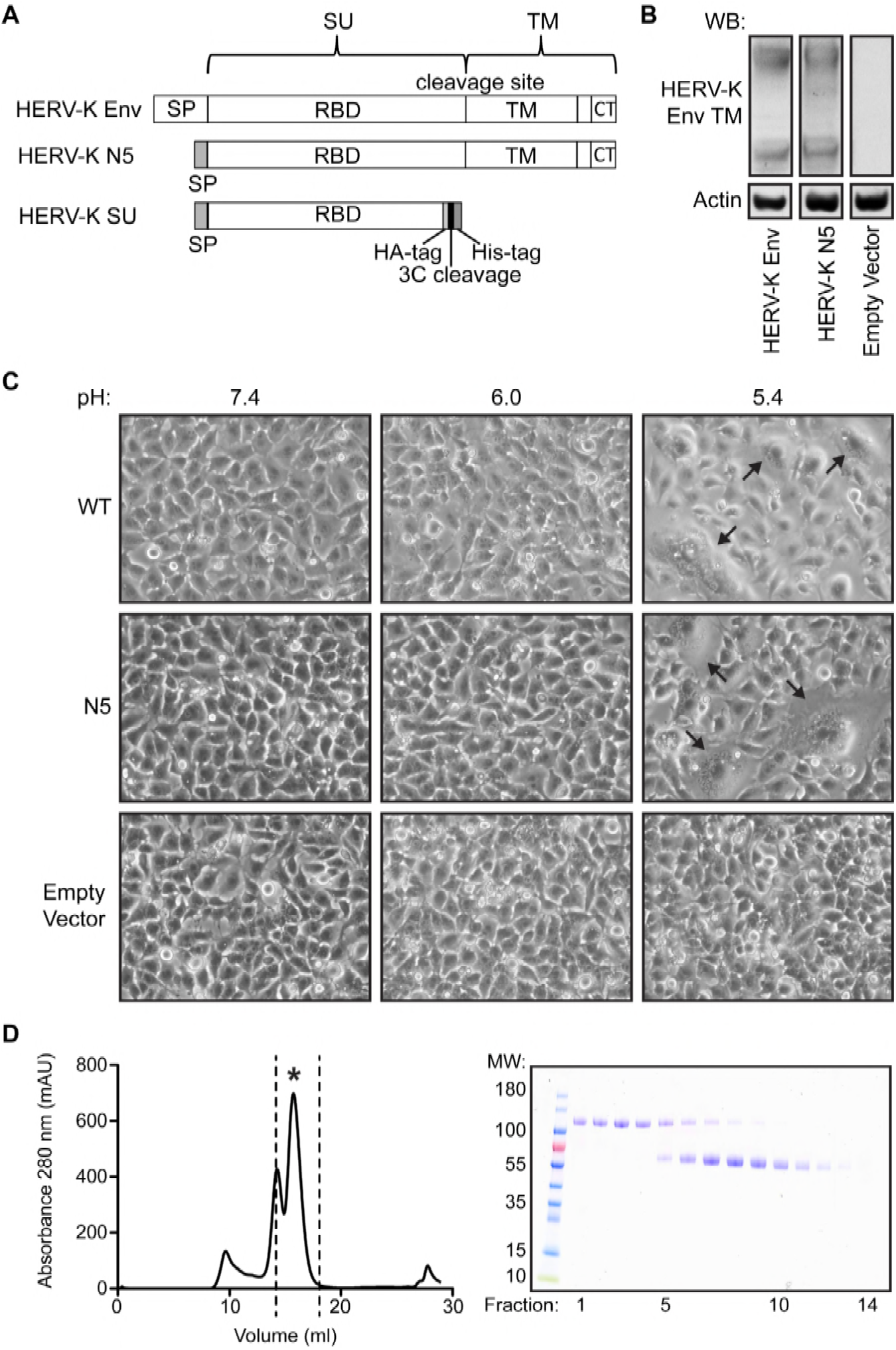
Validation of HERV-K SU. (A) Schematic of HERV-K Env, HERV-K N-terminal truncation N5, and HERV-K SU used in this study. **(B)** HERV-K Env and HERV-K N5 were transfected into BSRT7 cells and cell lysates were subjected to Western blot against HERV-K Env TM subunit and actin, to assess expression and proteolytic processing. For HERV-K Env blot: top band, uncleaved Env; bottom band, TM subunit. **(C)** BSRT7 cells were transfected with HERV-K Env, HERV-K N5, and empty vector. Cells were exposed to the indicated pH and assessed for the presence of multinucleated syncytia (indicated by arrows) **(D)** FPLC trace of HERV-K SU from gel filtration chromatography. The major peak (at approximately 15 ml, indicated with an asterisk) corresponds to monomeric SU. The peak at 13 ml corresponds to dimeric SU, and the peak at 9 ml is an aggregate of SU. Fractions from the FPLC (indicated with dashed lines) were run on a non-reducing SDS-PAGE and coomassie stained. Fraction 1 corresponds to 14.16 ml and fraction 14 corresponds to 18.06 ml. The top band at approximately 120 kDa represents the dimeric species and the lower band at approximately 60kDa represents monomer. Only fractions containing only monomer were used for pull-down experiments.

**Fig S7.**
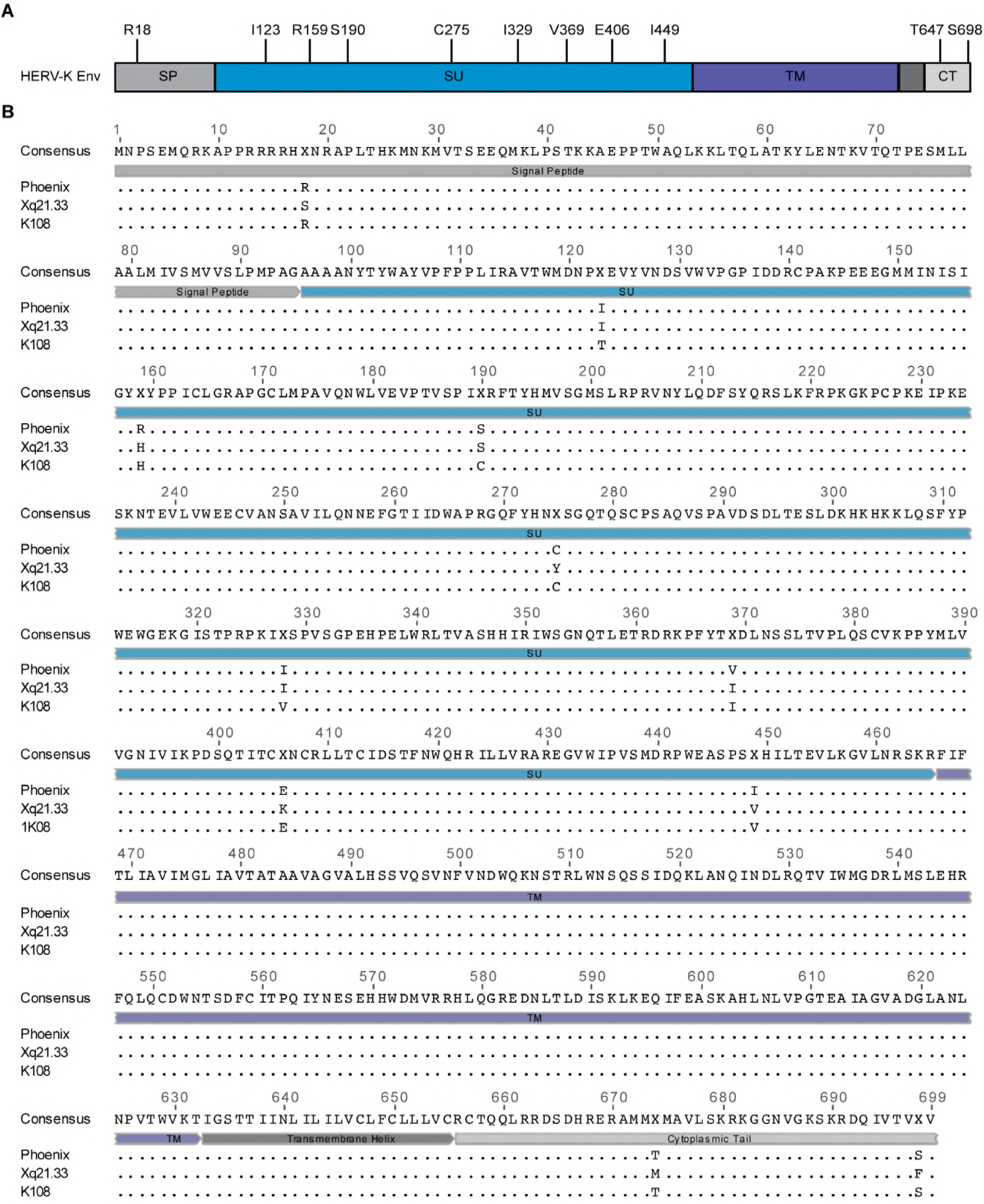
Alignment of Env sequences from Phoenix, Xq21.33, and HERV-K 108. (A) Schematic of HERV-K Env. Positions of amino acids with differences between Phoenix and either Xq21.33 or K108 are shown with the amino acid identity in Phoenix indicated. SP: signal peptide. SU: surface subunit. TM: transmembrane subunit. CT: cytoplasmic tail. **(B)** Alignment of Phoenix, Xq21.33, and K108 Envs.

**Fig S8.**
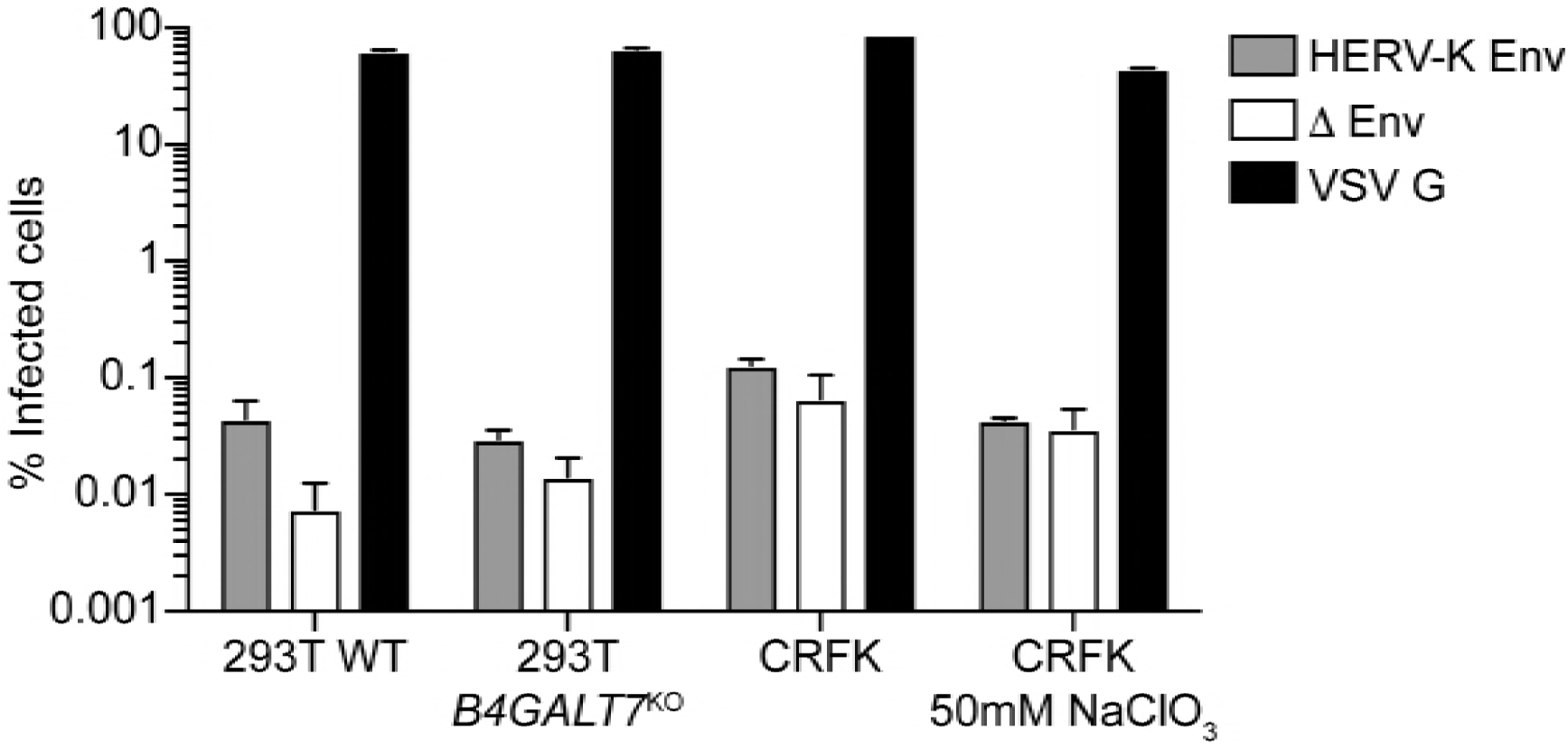
Relative infectivity of lentiviral pseudotypes. Lentivirus was produced as described above and particle concentration determined by p24 ELISA. The indicated cell lines were inoculated with equal particle numbers, based on p24 levels. % infected cells was determined by flow cytometry to determine the % GFP positive cells. Pseudotypes bearing HERV-K Env have an approximately 4-log defect in relative infectivity compared to those bearing VSV G, and have relative infectivities close to that of “bald” (Δ Env) particles.

